# Splice-specific deficiency of the PTSD-associated gene PAC1 leads to a paradoxical age-dependent stress behavior

**DOI:** 10.1101/2020.01.24.918185

**Authors:** Jakob Biran, Michael Gliksberg, Ido Shirat, Amrutha Swaminathan, Talia Levitas-Djerbi, Lior Appelbaum, Gil Levkowitz

## Abstract

The pituitary adenylate cyclase-activating polypeptide receptor (PAC1, also known as ADCYAP1R1) is associated with post-traumatic stress disorder and modulation of stress response in general. Alternative splicing of PAC1 results in multiple gene products, which differ in their mode of signalling and tissue distribution. However, the roles of distinct splice variants in the regulation of stress behavior is poorly understood. Alternative splicing of a short exon, which is known as the “hop cassette”, occurs during brain development and in response to stressful challenges. To examine the function of this variant, we generated a splice-specific zebrafish mutant lacking the hop cassette, which we designated ‘*hopless’*. We show that *hopless* mutant larvae display increased anxiety-like behavior, including reduced dark exploration and impaired habituation to dark exposure. Conversely, adult *hopless* mutants displayed superior ability to rebound from an acute stressor, as they exhibited reduced anxiety-like responses to an ensuing novelty stress. We propose that the developmental loss of a specific PAC1 splice variant mimics prolonged mild stress exposure, which in the long term, predisposes the organism’s stress response towards a resilient phenotype. Our study presents a unique genetic model demonstrating how early-life state of anxiety paradoxically correlates with reduced stress susceptibility in adulthood.

## Introduction

PAC1 (a.k.a. Adcyap1r1) is a G-protein coupled receptor (GPCR) that serves as the high-affinity receptor for pituitary adenylate cyclase-activating polypeptide (PACAP). PAC1 has pleiotropic functions and was demonstrated to be involved in the regulation of several homeostatic processes including metabolic rate and food consumption^1,2^, circadian rhythm^3^ and, in particular, stress response^4,5^. Intracerebroventricular injection of PACAP increased phosphorylated cyclic AMP response element binding protein (pCREB) and CRH immunoreactivity in the rat paraventricular nucleus^6^. PACAP knockout mice display blunted hypothalamic CRH levels in response to restraint challenge^7^. PACAP/PAC1 signaling was also associated with hypothalamo-pituitary-adrenal activity and stress-related behaviors in humans and rodents^5,8,9^. Furthermore, this pathway was correlated with stress-related risky behaviors in human and rodents^10,11^. Overall, these findings support positive stress regulation by PAC1; yet, some data suggest that it may also act to suppress stress phenotypes^4,12^.

It has been suggested that genetic vulnerability to post-traumatic stress disorder (PTSD) may depend on PAC1 expression and single-nucleotide polymorphism (SNP) in the PAC1 gene. Ressler et al. demonstrated that a specific PAC1 genotype is strongly correlated with susceptibility to PTSD in women, probably due to perturbed expression of PAC1 resulting in impaired stress responses^13^. The same PAC1 SNP was associated with PTSD in African-American females, emotional numbing in traumatized earthquake Chinese survivors, dark-enhanced startle response in children^14–16^, and with impaired hippocampal and amygdalar activation in response to fearful stimuli and contextual fear conditioning in non-traumatized individuals^17,18^. Moreover, PTSD was correlated with altered expression of PAC1 in the human cortex^13^.

These accumulating data have associated PAC1 to PTSD based on SNPs in the PAC1 gene promoter, global PAC1 expression or both. However, PAC1 mRNA undergoes extensive alternative splicing, resulting in the generation of several protein isoforms that differ in their ligand affinity and signal transduction cascades^19–21^. There are at least 17 known PAC1 splice isoforms. The predominant brain isoforms are PAC1-hop, which contains an alternatively spliced short exon termed the “hop cassette”, and PAC1-short that lacks it^22–24^. PAC1-hop encodes 28 amino acids of the third intracellular loop of this GPCR. PAC1-hop is expressed in neuroendocrine cells in mammals and it is widely expressed in the brain and gonad tissues^21,22,25^. The splicing of the hop cassette occurs during brain development and in response to stressful challenges. However, the specific roles of distinct splice variants in the regulation of stress-related behaviors are yet to be uncovered.

Neuroendocrine systems mediating homeostatic response to stressful challenges are well conserved between mammals and fish; indeed, PAC1 splice variants have been identified in zebrafish^21,26^. The zebrafish genome contains two PAC1 paralogs, namely *pac1a* and *pac1b;* however, only the *pac1a* gene contains the hop exon. We have previously demonstrated that alternative splicing of PAC1 is induced by an acute stress challenge to mice, which suggested that the ratio between PAC1-hop and PAC1-null isoforms may be involved in the adaptive response to stress^4^. We further demonstrated that zebrafish larvae injected with an antisense oligonucleotide that prevents the inclusion of the hop cassette displayed prolonged increase of *corticotropin-releasing hormone* (*crh*) transcription and impaired anxiety-like dark avoidance behavior^4^. In the present study, we examined the short- and long-term behavioral consequences of this splice-specific PAC1 deficiency by generating a germline-transmitted zebrafish mutant lacking the hop cassette, which we designated ‘*hopless’*. We show that while *hopless* larvae display increased anxiety-like and stress related behaviors, adult *hopless* mutants exhibit increased resilience to acute stress. These findings suggest that PAC1-hop splice isoform is involved in the developmental establishment of stress responsiveness.

## Results

### Generation of zebrafish mutant with germline deletion of the hop-coding exon

As mentioned, while PAC1 is known to be involved in modulation of stress response, the functions of each of its many splice variants are poorly understood. To examine the specific role of PAC1-hop splice variant, we have generated a zebrafish mutant that harbors a deletion of the alternatively-spliced hop exon of *pac1a* gene (**Fig. 1A**). To generate this mutant, we used the CRISPR/Cas9 system to simultaneously target two intragenic sites around exon 10 of *pac1a* (**Fig. 1A**). To avoid introduction of a phantom exon due to partial intronic retention, gRNA binding sites were designed at least 600 bp upstream and downstream to the hop cassette, thus including its predicted cis-regulatory splice sites. Mosaic F_0_ fish identified for carrying large genomic deletions were outbred with *Tüpfel long fin* (*TL*) wild-type (WT) strain. Their offspring were screened for individuals carrying at least 1,600 bp deletion in their genomic DNA. Four individual F_1_ fish were identified as carriers of the same genomic ∼1,600 bp indel. The resulting germline-transmitting mutant allele lacked the *hop* exon (**Fig. 1B**) as did the *pac1a* mRNA gene product (**Fig. 1C**); hence, we named this mutant ‘*hopless*’. Sequencing of *pac1a* cDNA that was derived from the *hopless* mutant line confirmed the in-frame deletion of exon 10 in the mature mRNA gene product, without perturbing the amino acid reading frame of the flanking exons 9 and 11.

**Figure 1.**
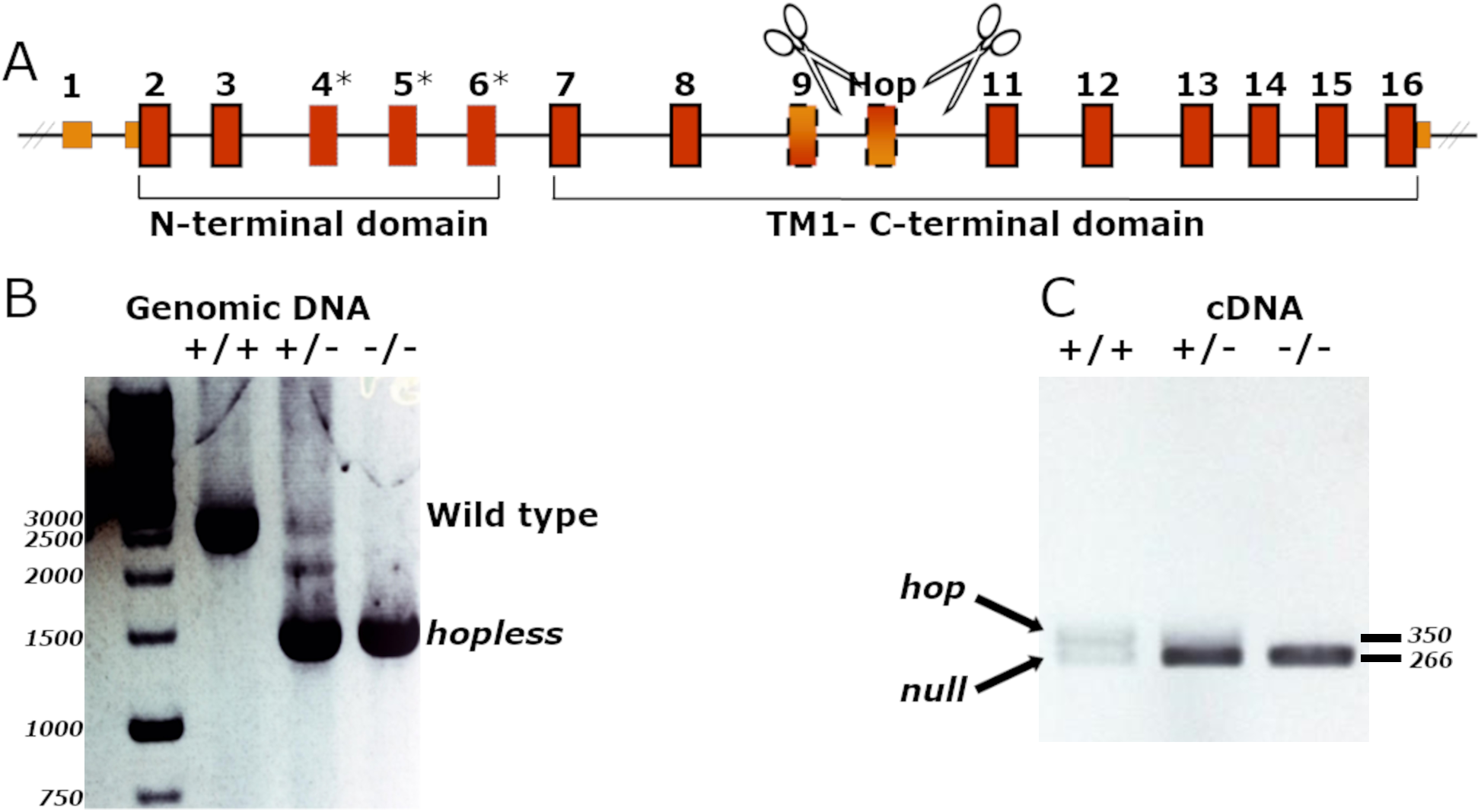
Establishment of *hopless^wz18^* zebrafish mutant. A pair of gRNAs (indicated by scissors) were designed to target the upstream and downstream genomic regions flanking the zebrafish hop cassette (**A**). PCR amplification of the genomic hop-cassette site demonstrate a single truncated product in the mutant genome (**B**). Amplification of PAC1-hop region in the zebrafish PAC1 mRNA shows loss of the hop isoform in cDNA produced from 6 dpf larvae (**C**). Plus sign represents wild-type allele; minus sign indicates *hopless* allele. Source images for the DNA gels electrophoresis are provided in Supplementary Figure 1.

Finally, homozygous *hopless* fish were viable and fertile and displayed no detectable morphological abnormalities compared to their WT siblings. Therefore, the generated mutant allowed us to assess the behavioral effects of the deleted *hop* exon in both larval and adult stages.

### *hopless* mutant larvae display heightened anxiety-like behaviors

To measure the behavioral effect of *hop* exon deficiency, we first used the light-dark preference test (**Fig. 2A**). In this assay, a larvae that is placed in a half lit, half dark arena displays a strong preference to the lit region^4,27^. The time spent in the dark is increased following exposure to anxiolytic drugs and, therefore, this behavioral assay is considered a measure of larval anxiety-like state^4,27^. We performed the test on 6 days post-fertilization (dpf) *hopless* larvae and genetically matched WT larvae. As previously demonstrated for larvae exposed to the anxiogenic caffeine^27^, we found that *hopless* larvae displayed significant reduction in the time spent in the dark zone, as well as in the number of entries to the dark zone (**Fig. 2B,C**). The observed phenotype was not due to locomotion defects, as the distance and speed of swimming of *hopless* larvae was similar to those of WT larvae throughout the test (**Fig. 2D,E**).

**Figure 2.**
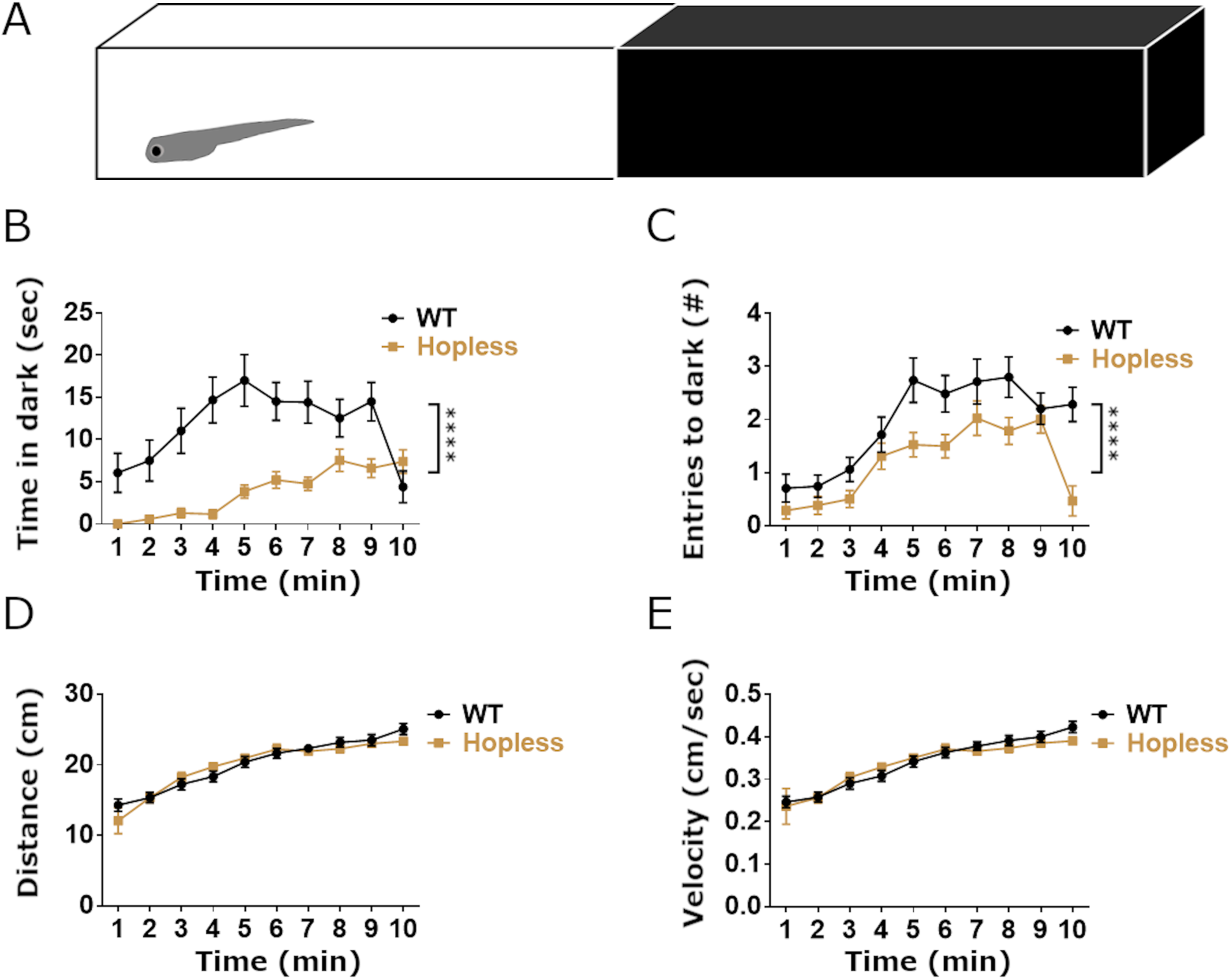
*Hopless* larvae display increased light preference in light-dark preference test. Larvae were placed on the lit side of a half lit-half dark rectangular arena and their behavior was recorded during 10 minutes (A). Analysis of locomotor activity showed that *hopless* mutants covered significantly less distance in the dark zone (**B**) and exhibited less entries into the dark (**C**) relative to WT larvae. Nonetheless, 6 dpf *hopless* larvae (n=42) did not differ from WT controls (n=35) in total distance swam (**D**) or average velocity (**E**). Data are presented as mean±SEM of 1 minute time bins. Data were analyzed using 2-way ANOVA. For all analyzed parameters, a significant effect of time (p<0.0001) was observed. Quadruple asterisks indicate significant difference between genotypes (p<0.0001). No significant interaction was found for time*genotype.

We next performed the light-dark transition assay, which measures the locomotion response of larvae to a sequence of sudden transitions between bright light and total darkness. Larvae tend to respond to sudden darkness by a short burst of vigorous swimming, the intensity and duration of which are increased under stressful conditions^28^. We analyzed larval locomotion using a 30 minute dark/light alternating periods^29^, as we found that this interval enables complete rebound from the stressful transitions (**Fig. 3A**). Results showed that immediately after light-to-dark transitions, WT larvae displayed darting behavior followed by a gradual decrease in locomotion, presumably due to acclimatization to the darkened environment (**Fig. 3A**). Conversely, dark-to-light transitions induced a short freezing episode, which was followed by a locomotion rebound (**Fig. 3A**). Moreover, the intensity of larval locomotion response to the above transitions was reduced during repeated light-dark cycles, indicating that they habituated to the sudden changes in the environment. To test whether the light-to-dark transition measures anxiety-like response, WT larvae were treated with diazepam, which was shown to be anxiolytic in both larvae and adult zebrafish^27,30^. Diazepam-treated larvae displayed reduced darting activity during the transition from light to dark, suggesting that light-to-dark sensory challenge is anxiogenic (**Fig. 3F,G**).

**Figure 3.**
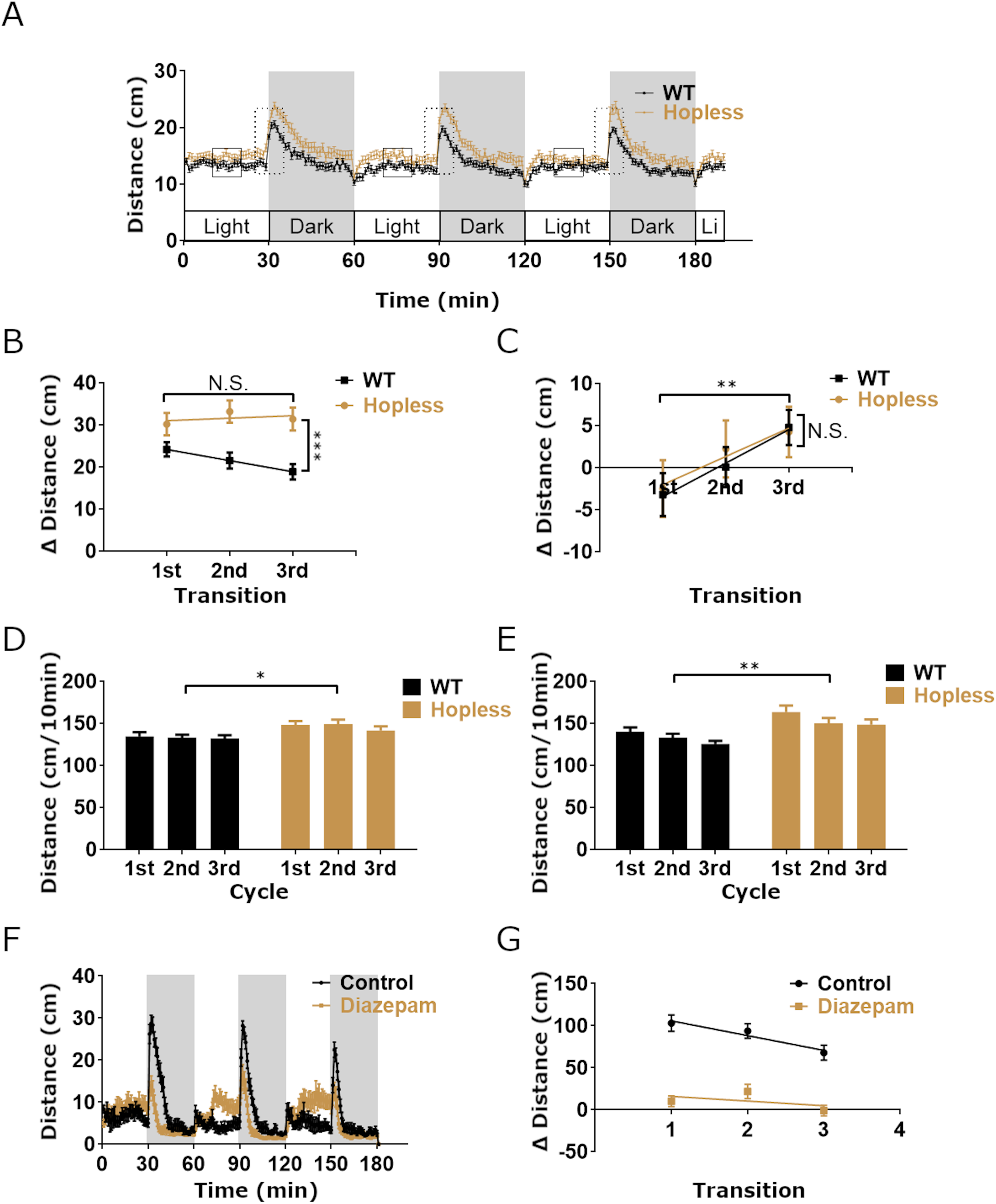
*Hopless* larvae display impaired stress-related behavior in light-dark transitions assay. Mean total locomotor activity of 6 dpf wild type (WT; n=48) and *hopless* mutant (n=41) larvae was measured during 3 h of 30 min alternating light-dark cycles (**A**). Mean total activity of each genotype was calculated for the middle 10 min interval of each phase (black squares in **A**) or as the difference in activity between 5 min post-transition and 5 min pre-transition (dashed rectangles in **A**). Analysis of larval activity during light-to-dark transitions revealed that WT larvae gradually decreased their dark-enhanced activity with each cycle, representing habituation to the challenge, whereas *hopless* larvae failed to display such habituation (**B**). During the opposite dark-to-light transitions no differential responses between genotypes were observed (**C**). In general, *hopless* larvae exhibited significantly increased activity under both light and dark conditions (**D** and **E**, respectively). WT larvae (n=24) that were treated with 5 µM diazepam displayed similar activity pattern to non-treated siblings (n=24; **F**); yet, treated larvae displayed reduced dark-enhanced activity (**G**). Data were analyzed using 2-way ANOVA. Single asterisks indicate p<0.05; double asterisks indicate p<0.01; triple asterisks indicate p<0.001. No significant interaction was found for time*genotype.

In contrast to the behavioral pattern observed in WT fish, *hopless* larvae failed to habituate to repeated light-to-dark transitions, as their darting response did not change between the first and third transitions (**Fig. 3B**). Interestingly, their habituation to sudden dark-to-light transitions was similar to that of the control, suggesting that this challenge may be less stressful to the larvae (**Fig. 3C**). Finally, we observed that *hopless* larvae exhibited heightened locomotion in both light and dark phases. This may indicate an impaired response of this mutant to the novel environment of the behavioral arena (**Fig. 3D,E**).

Taken together, our results indicate that *hopless* mutant larvae, which lack the alternatively spliced *hop* exon, display heightened anxiety-like responses to sensory-based light-dark stress stimuli.

### Adult *hopless* zebrafish display impaired behavioral response to novel environment

To assess whether the stress-related behavioral phenotypes of the *hopless* mutant persist to adulthood, we utilized the novel tank diving and open field assays. When adult zebrafish are introduced to a novel tank, they initially dive to the bottom half of the tank and, over time, they gradually begin to explore its top half^31^. This stereotypical behavior is commonly used to measure stress and anxiety-like response of zebrafish to a novel environment^32,33^. When we introduced adult WT or *hopless* zebrafish to the novel tank arena with no prior stressful challenge, both genotypes displayed similar behaviors, as reflected by similar swimming velocity and distance (Fig. 4A,B), as well as by the similar number of entries and time spent in the top half of the tank (**Fig. 4C,D**). Likewise, no significant differences between WT and *hopless* zebrafish were found in open field behavioral parameters, including locomotion, place preference towards center versus the walls, and freezing, which are also considered anxiety-like responses (**Supp. Fig. 2**). These results indicate that naïve adult *hopless* mutants do not display differential response in novelty-based behavioral assays.

**Figure 4.**
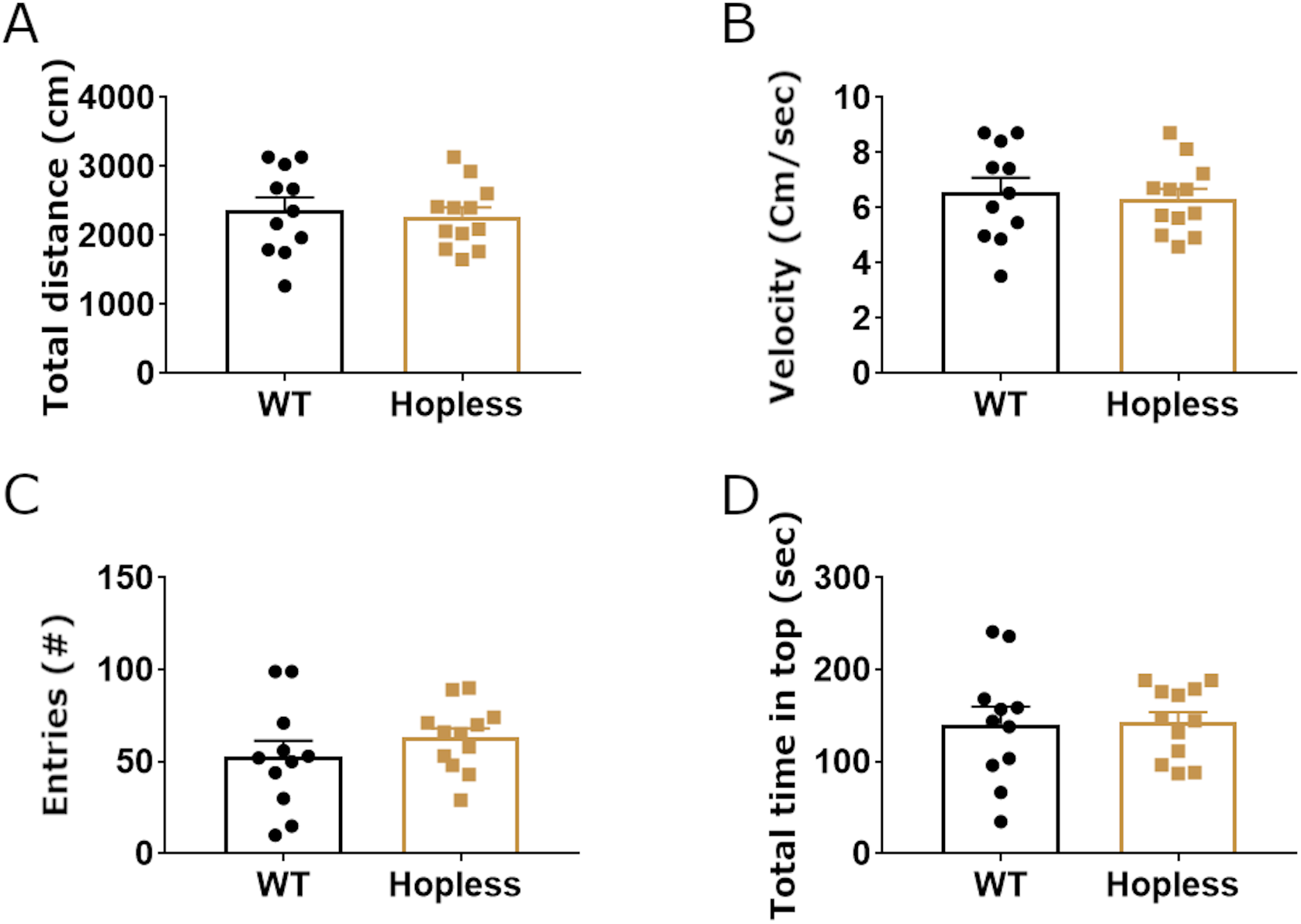
Naïve *hopless* adults display normal behavior in novel tank diving assay. The behavioral response of adult *hopless* mutants (n=12) and WT siblings (n=11) to a novel tank was measured. The two genotypes exhibited a similar activity level, as seen in their total swimming distance swum and average swimming velocity during the 10 min test (**A** and **B**). Both genotypes displayed similar exploration of the top half of the tank (**C** and **D**). Data was analyzed using students-t test.

We next examined the behavioral responses of *hopless* adults in novelty-based assays following an acute stressful challenge, namely a short restraint in a confined space. Wild-type and *hopless* siblings were restrained for 2 minutes in a 50 ml conical tube containing 5 ml of water, such that the fish was covered with water allowing normal breathing, but in a highly confined space. Thereafter, both genotypes were analyzed in the novel tank diving and open field assays. Novel tank diving assay duration was extended to 14 minutes, in order to allow the restrained fish to recover from the stressful restraint challenge.

When placed in the novel tank arena, stressed fish of both genotypes covered the same distance with a similar swim velocity (**Fig. 5A,B**). However, *hopless* mutants displayed more entries to the top half of the tank, indicative of reduced stress behavior (**Fig. 5C**). *hopless* mutants also spent more time and covered more distance in the top half of the arena (**Fig. 5D-F**). When analyzed in the open field arena, stressed *hopless* mutants covered more distance with higher average velocity than stressed WT zebrafish (**Fig. 6A,B**). This effect was probably due to the reduced freezing of *hopless* mutants (**Fig. 6C**). Although no significant difference was found in the time spent in the center of the arena, *hopless* mutants performed significantly more visits to the center and showed a tendency to spend less time per visit, indicative of increased arena exploration (**Fig. 6D-F**). It is noteworthy that following restraint, WT fish froze also in the center of the arena (data not shown), which may explain, at least in part, the lack of difference in time spent in the center. We conclude that adult *hopless* mutants display improved behavioral recovery following an acute stress, suggesting that they are more resilient to stressful challenges.

**Figure 5.**
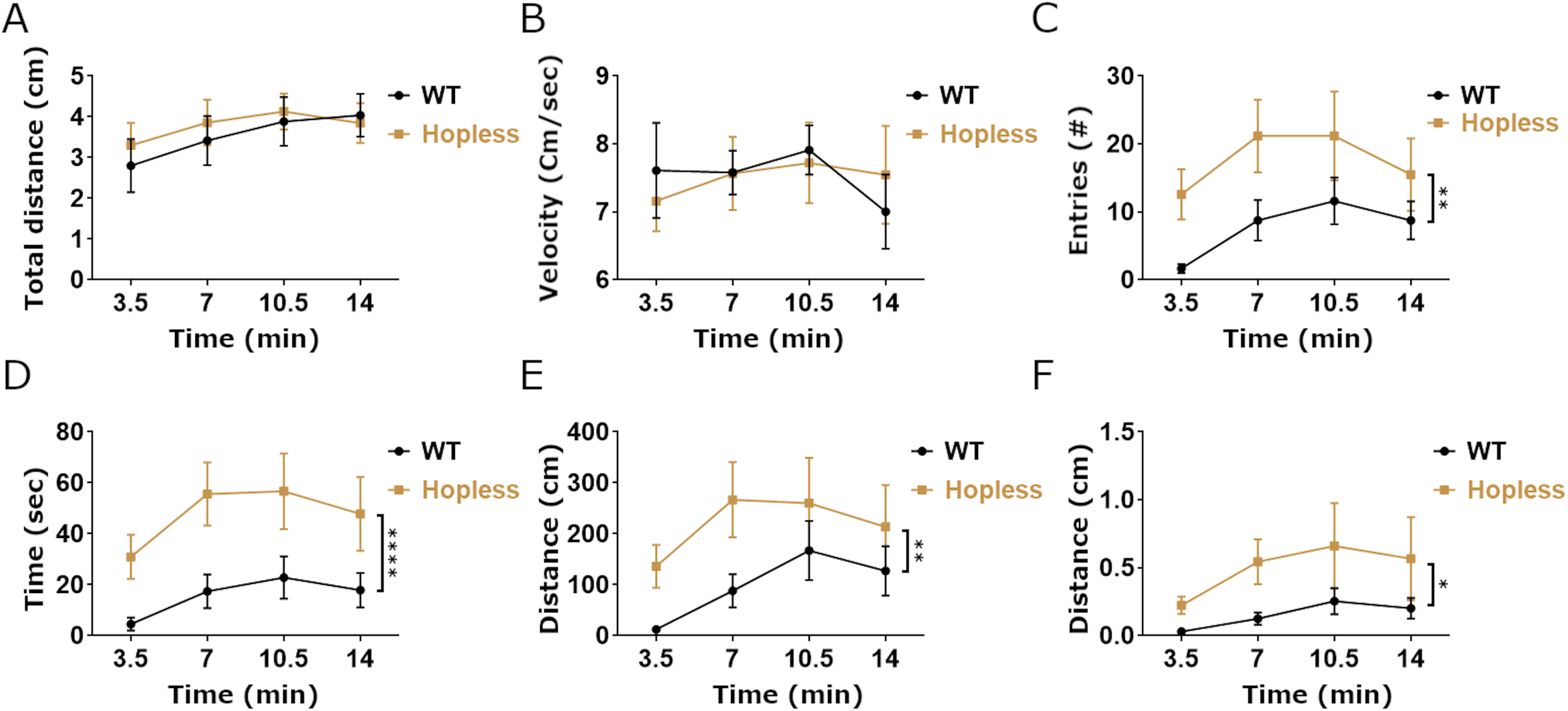
A preceding stressful challenge leads to blunted stress-related behavior of *hopless* adults in the novel tank assay. The behavioral response of adult *hopless* mutants (n=11) and WT siblings (n=13) to a novel tank was measured following 2 min restraint in a confined space. The two genotypes exhibited similar activity, as seen in their mean swimming distance and velocity (**A** and **B**). Nonetheless, when subjected to a preceding restraint stressfull challenge we observed significantly different behaviors between genotypes, namely that *hopless* mutants entered more times to the top half of the tank, spent more time and covered more distance there, even when normalized to distance swum at the bottom of the tank (**C**-**F**). Data were analyzed using 2-way ANOVA. Single asterisks indicate p<0.05; double asterisks indicate p<0.01; quadruple asterisks indicate p<0.0001.

**Figure 6.**
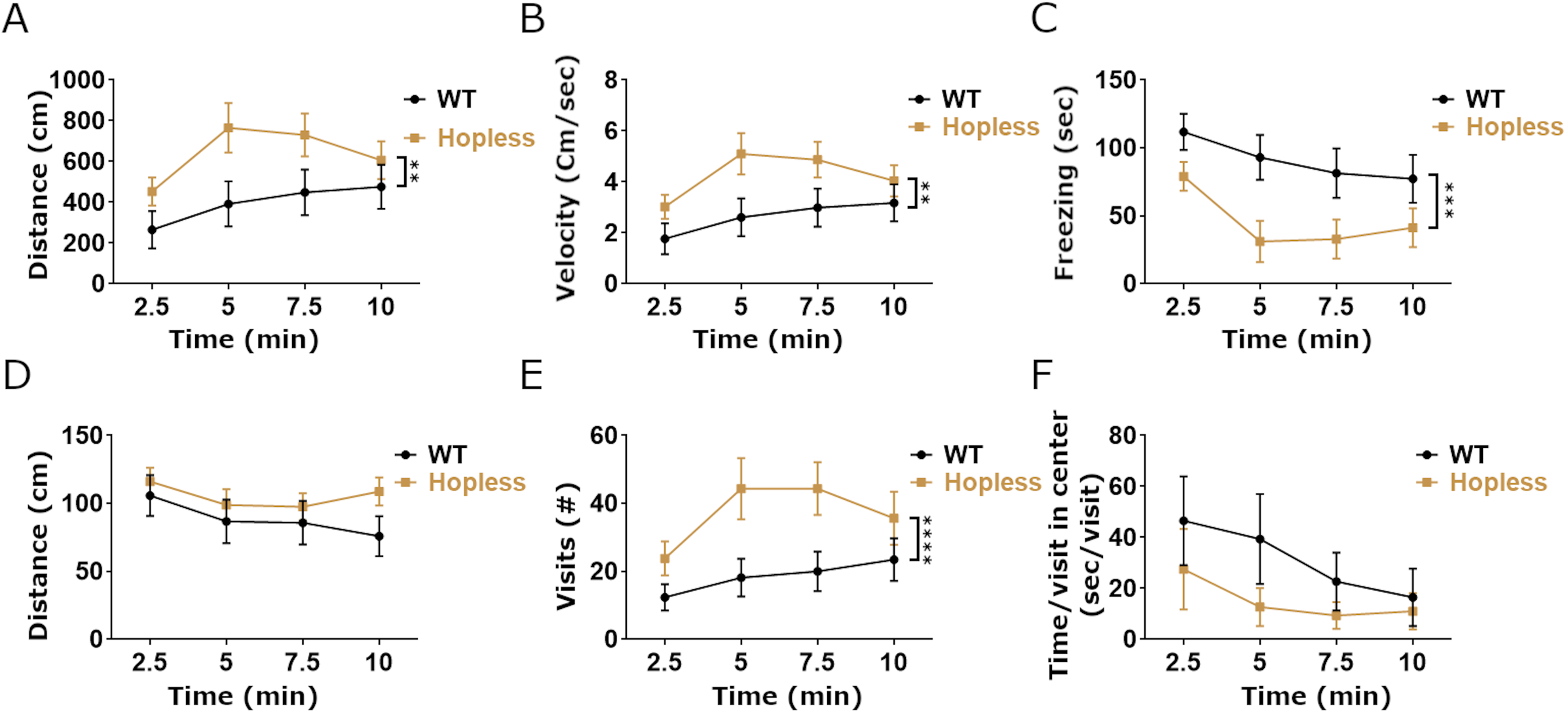
A preceding stressful challenge leads to blunted stress-related behavior of adult *hopless* zebrafish in the open field assay. Following 2 min restraint in a confined space, adult *hopless* mutants (n=10) and WT siblings (n=13) behavior was measured in a novel open field arena. A significant reduction is seen in activity level of WT, but not of *hopless* mutant zebrafish (**A** and **B**). This effect was probably due to the increased freezing of WT adults (**C**). No significant difference was seen in the time spent in the center of the open field arena (**D**); however, *hopless* mutants displayed significantly increased number of visits to the arena center and a tendency (p=0.09) to spend less time in each visit to center (**E** and **F**). Data were analyzed using 2-way ANOVA. Double asterisks indicate p<0.01; triple asterisks indicate p<0.001; quadruple asterisks indicate p<0.0001.

## Discussion

PAC1 is an important player in the regulation of physiological and behavioral stress responses. Perturbation of its expression was associated with PTSD and impaired stress responses^34,35^. Yet, although PAC1 undergoes extensive alternative splicing, the specific roles of its isoforms in the regulation of stress response have not been elucidated. In the present study, we have generated a unique genetic model, which lacks the *hop* exon, and used this mutant to examine the role of PAC1-hop splice variant in the regulation of stress-related behaviors throughout life. Our results revealed an intriguing phenomenon of an age-dependent switch in stress-related behaviors. As larvae, *hopless* mutants display increased anxiety-like responses. Conversely, adult *hopless* fish rebounded better from a major stressor (restraint), displaying reduced anxiety-like responses to a subsequent milder stressor (introduction into novel environment), which indicates improved resilience to stress during adulthood. We suggest that the developmental loss of a specific PAC1 splice variant mimics a heightened anxiety state that, in the long-term, renders an improved ability to cope with stressful challenges.

PAC1 splice isoforms are known to mediate distinct signal transduction pathways due to differential coupling to G-proteins^21,36,37^. The PAC1-null isoform, which does not contain the hop exon, displays very low or no activation of PLCβ, whereas PAC1-hop1 switches its mode of signaling from AC to PLCβ^38–42^. PAC1-hop1 was also shown to regulate Ca^2+^ mobilization and neurosecretion^43–46^. Such differential signal transduction could affect behavioral phenotypes, as previously demonstrated for other GPCRs^47–50^. Accordingly, we have previously shown in adult mice that acute foot-shock stressor induces a change in the ratio between PAC1-hop and PAC1-null splice isoforms in the hypothalamic paraventricular nucleus^4^.

Using an antisense morpholino oligonucleotide in larval fish, we also showed that interference with PAC1a-hop alternative splicing leads to impaired *crh* transcriptional dynamics, as well as adaptation to a stressful challenge^4^. However, due to the transient nature of this gene-knockdown approach, the long-term consequences of developmental deficiency of PAC1-hop on adult stress responses were not examined. In the present study, we utilized the CRISPR/Cas9 method to generate a germline mutation resulting in the loss of PAC1-hop isoform mRNA without affecting the generation of the other PAC1 isoforms, allowing functional investigation of PAC1-hop.

Early life stress exposure may have negative or positive impact on adult psychological well-being. For example, in humans early life stress is known to increase the risk for psychopathologies in adulthood^51,52^. In particular, PTSD was recently correlated with childhood anxiety of both military and civilian men^52^. Nonetheless, more than 60 years ago, Levine demonstrated that mild stress exposure can lead to an inoculation effect in the adult rat, making it more resilient to stressors^53^. These findings were later validated in mice and squirrel monkeys^54,55^. In line with this notion, in our study *hopless* larvae exhibited increased anxiety-like behavior in the light-dark preference test and reduced habituation to repeated light-to-dark transitions. However, pre-challenged adult mutants displayed a blunted stress response of increased exploration and reduced freezing in both novel tank diving and open field assays, indicating enhanced resilience to novelty stressors.

As mentioned above, in various animal models, PAC1 signaling has been mainly associated with activation of physiological and behavioral stress responses, including PTSD^10,13, 56–58^. Infusion of PACAP, the high-affinity PAC1 ligand, to the bed nucleus of stria terminalis (BNST) potentiated anxiogenic response in pre-stressed but not in naïve male rats, which was likely conveyed by PAC1 signaling^9,59^. PAC1-KO mice display blunted elevation in corticosterone following acute restraint stress, but not under naïve conditions^5^. However, our current study highlights the need for better understanding of the physiological and psychological functions of specific PAC1 splice isoforms. Generation of additional site- and splice-specific models is needed to further elucidate mechanisms underlying splicing-based regulation of the stress response.

Although the underlying mechanism for the switch in stress responsiveness of *hopless* mutants studied herein is still unknown, some studies may provide hints. A dynamic shift in the ratio of PAC1-hop and PAC1-short isoforms was shown to regulate pro- and anti-mitogenic activity of PACAP in the developing brain^60^. The ratio of PAC1-hop2 mRNA to other PAC1 variants in the ventromedial nucleus of the hypothalamus of female rats appears to be dynamic, depending on different steroid environments^61^. This study suggests that steroid-regulated changes in the expression of PAC1 splice variant in the hypothalamus contribute to the effect of PACAP on female receptivity^61,62^. We have previously suggested that alternative splicing of the hop cassette serves as an ON/OFF stress switch^21^, in line with our finding that stress-induced alternative splicing of PAC1-hop is involved in the termination of *crh* transcription and adaptive anxiety-like larval behavior. Larvae with depleted PAC1-hop1 display delayed dark-avoidance recovery following a stressful challenge response^4^.

Taken together, our findings suggest that PAC1-hop is involved in the developmental establishment of stress responsiveness. Thus, in our current study, *hopless* larvae likely experienced an impaired ability to terminate their stress response, which presumably led to prolonged and possibly heightened anxiety-like states during early life stages. These states could occur in response to any of the very mild stressful events that the animals experience during rearing in an enclosed laboratory setting, for example, the sudden daily switch from dark to light, encounter with human caretakers, tank cleaning, etc. We submit that in the case of the *hopless* mutant, this early life state of anxiety paradoxically correlates with reduced stress susceptibility in adulthood, reminiscent to the early-life stress inoculation proposed by Levin ^53^ and others.

## Materials and methods

### Animal care and maintenance

Zebrafish were bred and reared at 28.5°C under 14 h/10 h light/dark cycle, according to standard protocols. Embryos were raised at 28.5°C in 30% Danieau’s medium (0.17 mM NaCl, 0.21 mM KCl, 0.12 mM MgSO_4_, 0.18 mM Ca(NO_3_)_2_, 0.15 mM HEPES, pH 7.4) supplemented with 0.01 mg/L methylene blue. All experimental procedures were approved by and conducted in accordance with the Weizmann Institute’s Institutional Animal Care and Use Committee (IACUC).

### Genome editing using CRISPR/Cas9

To generate the *hopless^wz18^* allele (ZFIN ID: ZDB-ALT-191028-3) we employed genome editing as previously described ^63,64^, with slight modifications. Cas9 protein was produced by the Weizmann Institute of Science Protein Purification Unit using the pET-28b-Cas9-His (Alex Schier Lab Plasmids, Addgene, Cambridge, MA, United States) as a template. CRISPR sgRNAs (Supplementary Table S1) were designed using CHOPCHOP ^65^. Oligonucleotide containing the SP6 promoter sequence upstream of specific target sites was annealed with a constant oligonucleotide bearing Cas9 binding site. sgRNA were generated by *in vitro* transcription using a SP6 RNA polymerase MEGA script SP6 kit (Life Technologies, United States) and purified using miRNeasy kit (Qiagen, Germantown, MD, United States). Cas9 protein (600 ng) and sgRNAs (1500-2000 ng) were co-injected to *Tupfel long fin* (TL) zygotes at the one-cell stage. PCR analysis of ten embryos was performed to evaluate genomic mutations and occurrence of large deletions induced by multiple gRNAs injection. Injected siblings were raised to adulthood and screened for large deletions in genomic-DNA (gDNA) extracted from tail clips. Fish that were found to be positive for large genomic deletions were crossed with TL zebrafish to identify germline transmission. Germline heterozygotes were genotyped using Sanger sequencing and fish carrying the *hopless* allele were crossed to generate homozygous mutants.

### DNA extraction and genotyping

gDNA was extracted from tail fin clips of adult fish or whole embryo at 24 hours post-fertilization. DNA was extracted according to ^66^. Briefly, samples were digested at 50°C with 1% SDS and proteinase K solution (20 mg/ml). Protein precipitation was performed by adding 5 M NaCl solution and centrifugation. Supernatant was transferred to a clean tube, followed by DNA precipitation using ice-cold isopropanol. DNA pellet was isolated by centrifugation and washed with 70% ethanol. DNA pellets were air-dried, dissolved in 50 µl double distilled water and stored at −20°C until further analysis. The genomic region of *hop* cassette was amplified by using Taq DNA Polymerase Master Mix Red 2x reaction mix (Ampliqon, Odense, Denmark). Amplified DNA region was detected by 1-1.5% agarose gel electrophoresis, then extracted and purified by using the NucleoSpin® Gel and PCR Clean-up (Macherey-Nagel, Düren, Germany). Purified DNA fragments were sequenced by the Biological Services Unit at the Weizmann Institute of Science. Results were analyzed by ApE-A software (version 2.0, by M. Wayne Davis, University of Utah). See Supplementary Table S1 for oligonucleotide sequences.

### Light/dark preference test

Anxiety-like behavior of zebrafish larvae was measured by recording the preference of 6-day-old zebrafish larvae for either side of a custom-made light-dark arena. The test was generally performed according to ^4^ with slight modifications. Briefly, larval activity was recorded for a 10 min time period by high resolution infrared video camera (Flare 2M360-CL, I/O industries, London Ontario), with an image acquisition Sapera LT-development package (Teledyne Dalsa, Waterloo, Ontario) and recorded with Stream5 software (IO Industries, London, Ontario). The resulting videos were analyzed using EthoVision XT ((Noldus Information Technology, Wageningen, Netherlands). *Hopless* larvae and matched TL control larvae were randomly assigned into separate arenas with bottom lighting. Each arena was divided into two equal compartments in which the top, bottom and sides of the dark half were covered with Optical Cast Infrared (IR) Longpass Filters, allowing penetration of infrared light (Edmund Optics, Barrington, NJ, USA). The total distance swum in the whole arena and in each compartment was calculated and compared for each genotype. Animals that stopped swimming for periods of 60 s or more were removed from the analysis.

### Larval light/dark transitions test

Larval responses to sudden transitions from light to dark and from dark to light were monitored using the Noldus DanioVision tracking system (Noldus Information Technology, Wageningen, Netherlands), as previously described ^29^. For each experiment, individual larvae were placed in 48-well plates, which were put in the DanioVision system with light on for 1 h prior to the beginning of the trial. Larvae were subjected to three cycles of 30 min light/30 min dark followed by 10 min light. The experiment was repeated three times, each with a new batch of 6 dpf *hopless* and TL larvae. Total swimming distance and velocity were calculated using EthoVision XT Software (Noldus Information Technology). For the diazepam treatment experiment, larvae were exposed to 5 µM diazepam (Renaudin, France) for 2 h prior to the start of the behavioral testing. Control larvae were kept in similar conditions during this period. At the end of the incubation, larvae were transferred into individual wells of a 48-well plate and subjected to 30 min light/30 min dark cycles as described above.

### Open field

Behavioral phenotyping in open field arena was performed using a custom-made apparatus, as was previously described ^67^. Fish were placed in a circular arena of 23 cm in diameter filled to a height of 5 cm with regular system water. The arena was placed in a custom-built enclosure that restricted all visual cues from the laboratory frame, and was top-lit with ambient LED light. Videos were acquired with a 2M360-CL camera (IO Industries, London, Ontario), with an image acquisition Sapera LT-development package (Teledyne Dalsa, Waterloo, Ontario) and recorded with Stream5 software (IO Industries, London, Ontario). Tracking analysis was performed using EthoVision video tracking system (Noldus Information Technologies, Wageningen, Netherlands).

### Novel tank diving assay

Novel tank diving assay was performed as was previously described ^4^. The assay was conducted within the same enclosure as the open field assay and using the same recording equipment.

### Restraint challenge

Mild stress was induced in adult fish by placing individual fish in a standard 50 ml conical tube with 5 ml of system water for 2 min. The tube was placed in an upright position such that the fish could not swim, but was completely covered with water to prevent dehydration or loss of oxygen. Following the 2 min restraint, the fish were introduced to the open field arena or novel tank diving arena for behavioral tracking.

### Statistical analysis

Data are presented as mean ± standard error of the mean (SEM) and were analyzed using GraphPad Prism 7.01. All data sets were tested for departures from normality with Shapiro-Wilks test. Student’s *t*-test was used for all comparisons between two groups. 2-Way ANOVA was used for comparing multiple variables. Single asterisks indicates p<0.05, double asterisks indicates <0.01, triple asterisks indicates p<0.001 and quadruple asterisks indicates p<0.0001.

## Acknowledgements

We thank Roy Hofi for animal care; Nitzan Konstantin for English editing. G.L. is supported by the Israel Science Foundation (#1511/16); US-Israel Bi-National Science Foundation (#2017325); Yeda-Sela Center for Basic Research (in the frame of the Weizmann Institute) and the Nella and Leon Benoziyo Center for Neurological Diseases. G.L. is an incumbent of the Elias Sourasky Professorial Chair.

## Authors’ contributions

J.B. and G.L. planned the project. J.B. and M.G. and I.S. performed the behavioral experiments and analyzed the data. A.S. performed the diazepam experiment. L.A. and T.L.D. contributed to the initial establishment of the light-dark transition assay. J.B. prepared the figures. J.B. and G.L. wrote the manuscript. All authors reviewed the manuscript’s text.

## Supplementary Information

### Legends to Supplementary Figures

**Supplementary Figure 1.**
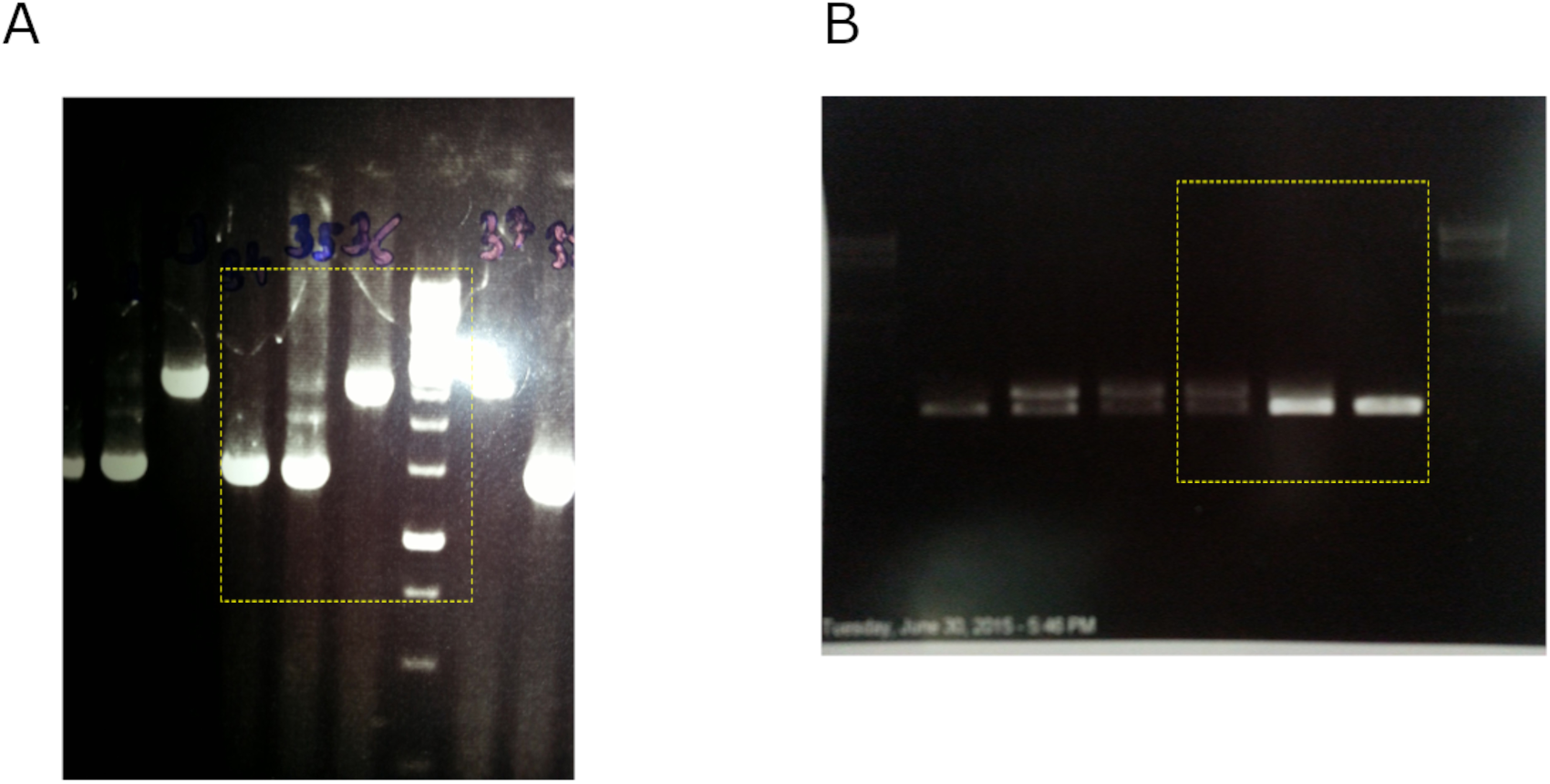
Source uncropped gel electrophoresis images of genomic DNA (**A**) and cDNA (**B**) in Figure 1. Images were cropped at the yellow dashed rectangle and black & white color was inverted for better contrast.

**Supplementary Figure 2.**
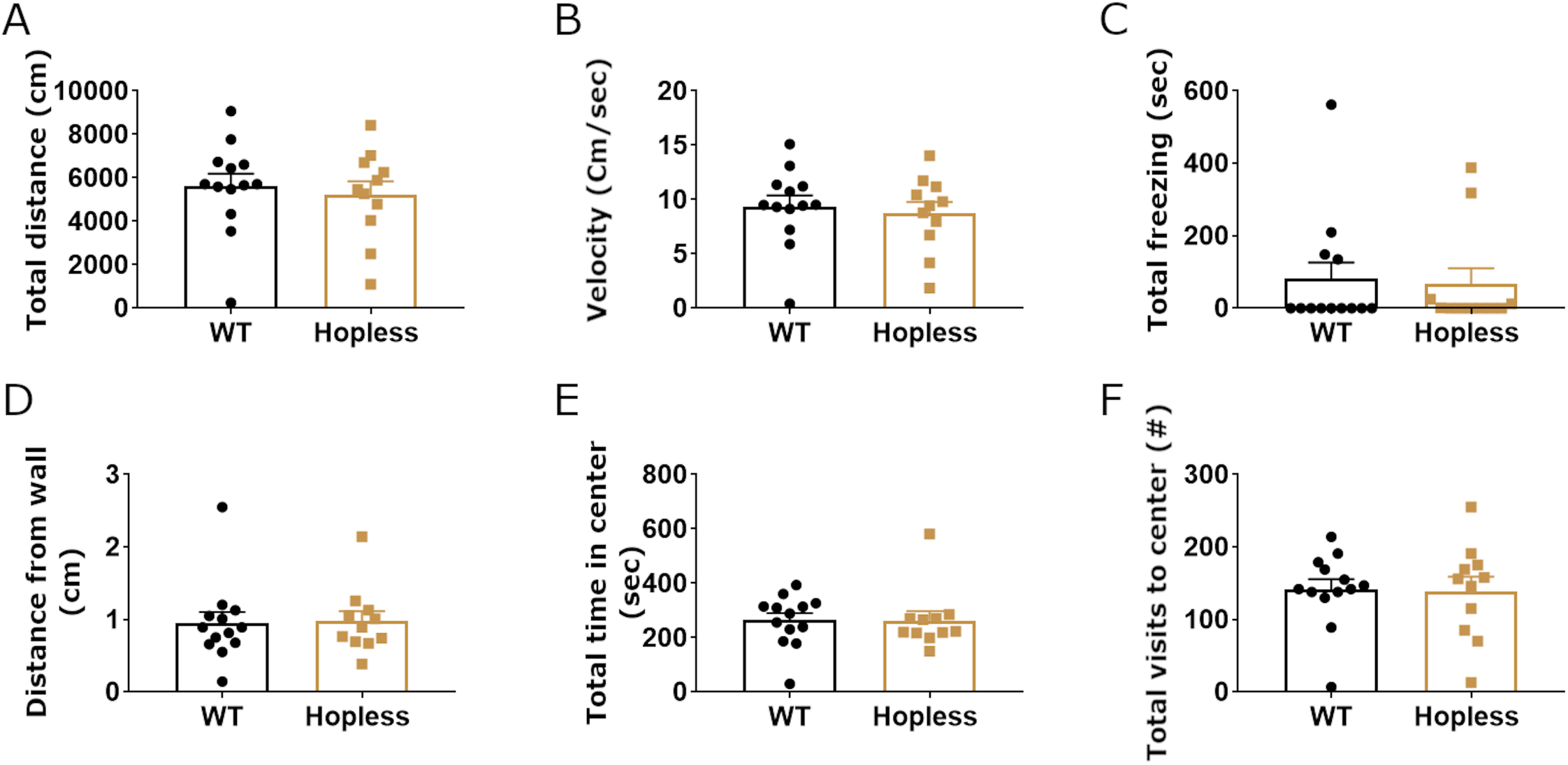
Naïve adult *hopless* mutants display normal behavior in the open field assay. The behavioral response of adult *hopless* mutants (n=11) and WT siblings (n=13) in a novel open field arena was measured. Similarly to the novel tank diving assay, the two genotypes exhibited a similar activity level, as seen in their mean swimming distance and velocity, which significantly increased with time (**A** and **B**). In addition, both genotypes displayed significantly decreased freezing, distance from walls and time spent in the center of the arena (**C**-**E**) and a slight but significant increase in their visits to the center (**F**), suggesting increased exploration activity over time. Data were analyzed using Student’s *t*-test.

**Supplementary Table 1.**
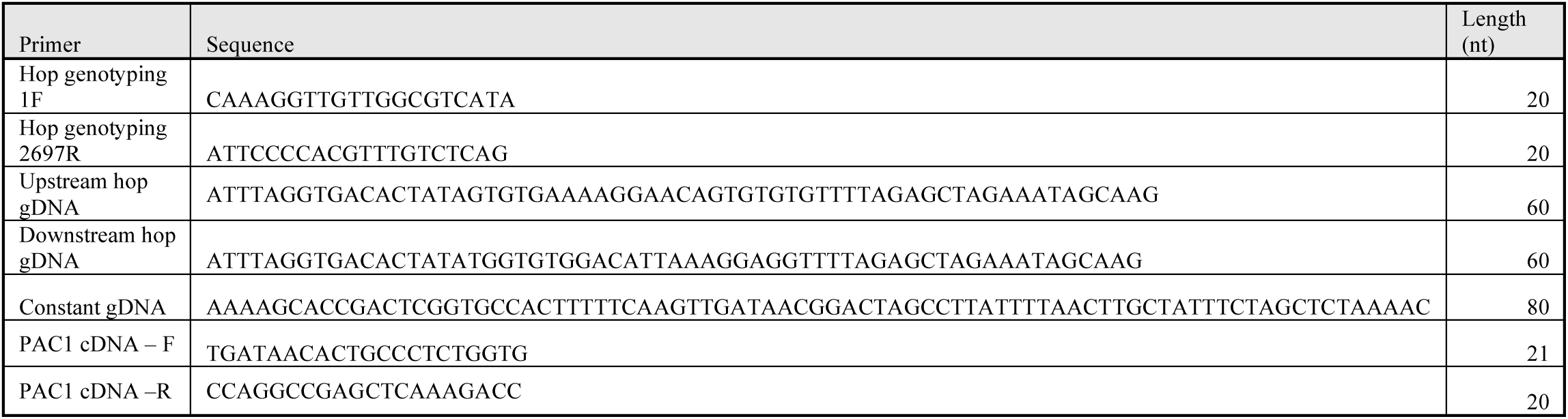
Oligonucleotides used for gRNA transcription and RT-PCRs.

## References

1 Huang, J., Waters, K. A. & Machaalani, R. Pituitary adenylate cyclase activating polypeptide (PACAP) and its receptor 1 (PAC1) in the human infant brain and changes in the Sudden Infant Death Syndrome (SIDS). Neurobiology of Disease 103, 70–77, doi:https://doi.org/10.1016/j.nbd.2017.04.002 (2017).

2 Mounien, L. et al. Pituitary Adenylate Cyclase-Activating Polypeptide Inhibits Food Intake in Mice Through Activation of the Hypothalamic Melanocortin System. Neuropsychopharmacology 34, 424, doi:10.1038/npp.2008.73 (2008).

3 Holland, P. R., Barloese, M. & Fahrenkrug, J. PACAP in hypothalamic regulation of sleep and circadian rhythm: importance for headache. The Journal of Headache and Pain 19, 20, doi:10.1186/s10194-018-0844-4 (2018).

4 Amir-Zilberstein, L. et al. Homeodomain protein otp and activity-dependent splicing modulate neuronal adaptation to stress. Neuron 73, 279–291, doi:10.1016/j.neuron.2011.11.019 (2012).

5 Mustafa, T. et al. Impact of PACAP and PAC1 receptor deficiency on the neurochemical and behavioral effects of acute and chronic restraint stress in male C57BL/6 mice. Stress 18, 408–418, doi:10.3109/10253890.2015.1025044 (2015).

6 Agarwal, A., Halvorson, L. M. & Legradi, G. Pituitary adenylate cyclase-activating polypeptide (PACAP) mimics neuroendocrine and behavioral manifestations of stress: Evidence for PKA-mediated expression of the corticotropin-releasing hormone (CRH) gene. Brain research. Molecular brain research 138, 45–57, doi:10.1016/j.molbrainres.2005.03.016 (2005).

7 Stroth, N. & Eiden, L. E. Stress hormone synthesis in mouse hypothalamus and adrenal gland triggered by restraint is dependent on pituitary adenylate cyclase-activating polypeptide signaling. Neuroscience 165, 1025–1030, doi:10.1016/j.neuroscience.2009.11.023 (2010).

8 Hammack, S. E. et al. Roles for pituitary adenylate cyclase-activating peptide (PACAP) expression and signaling in the bed nucleus of the stria terminalis (BNST) in mediating the behavioral consequences of chronic stress. Journal of molecular neuroscience : MN 42, 327–340, doi:10.1007/s12031-010-9364-7 (2010).

9 Roman, C. W. et al. PAC1 receptor antagonism in the bed nucleus of the stria terminalis (BNST) attenuates the endocrine and behavioral consequences of chronic stress. Psychoneuroendocrinology 47, 151–165, doi:10.1016/j.psyneuen.2014.05.014 (2014).

10 Dragan, W. Ł., Czerski, P. M. & Dragan, M. PAC1 receptor (ADCYAP1R1) genotype and problematic alcohol use in a sample of young women. Neuropsychiatric Disease and Treatment 13, 1483–1489, doi:10.2147/NDT.S137331 (2017).

11 Miles, O. W. et al. Pituitary Adenylate Cyclase-Activating Peptide in the Bed Nucleus of the Stria Terminalis Mediates Stress-Induced Reinstatement of Cocaine Seeking in Rats. Neuropsychopharmacology 43, 978, doi:10.1038/npp.2017.135 (2017).

12 Pilzer, I. & Gozes, I. VIP provides cellular protection through a specific splice variant of the PACAP receptor: A new neuroprotection target. Peptides 27, 2867–2876, doi:https://doi.org/10.1016/j.peptides.2006.06.007 (2006).

13 Ressler, K. J. et al. Post-traumatic stress disorder is associated with PACAP and the PAC1 receptor. Nature 470, 492–497, doi:10.1038/nature09856 (2011).

14 Wang, L. et al. PAC1 receptor (ADCYAP1R1) genotype is associated with PTSD’s emotional numbing symptoms in Chinese earthquake survivors. Journal of Affective Disorders 150, 156–159, doi:https://doi.org/10.1016/j.jad.2013.01.010 (2013).

15 Jovanovic, T. et al. PAC1 receptor (ADCYAP1R1) genotype is associated with dark-enhanced startle in children. Molecular Psychiatry 18, 742, doi:10.1038/mp.2012.98 (2012).

16 Almli, L. M. et al. ADCYAP1R1 genotype associates with post-traumatic stress symptoms in highly traumatized African-American females. American Journal of Medical Genetics Part B: Neuropsychiatric Genetics 162, 262–272, doi:10.1002/ajmg.b.32145 (2013).

17 Pohlack, S. T. et al. Neural Mechanism of a Sex-Specific Risk Variant for Posttraumatic Stress Disorder in the Type I Receptor of the Pituitary Adenylate Cyclase Activating Polypeptide. Biological Psychiatry 78, 840–847, doi:https://doi.org/10.1016/j.biopsych.2014.12.018 (2015).

18 Stevens, J. S. et al. PACAP receptor gene polymorphism impacts fear responses in the amygdala and hippocampus. Proceedings of the National Academy of Sciences 111, 3158, doi:10.1073/pnas.1318954111 (2014).

19 Holighaus, Y., Mustafa, T. & Eiden, L. E. PAC1hop, null and hip receptors mediate differential signaling through cyclic AMP and calcium leading to splice variant-specific gene induction in neural cells. Peptides 32, 1647–1655, doi:10.1016/j.peptides.2011.06.004 (2011).

20 Ushiyama, M. et al. Differential Intracellular Signaling through PAC1 Isoforms as a Result of Alternative Splicing in the First Extracellular Domain and the Third Intracellular Loop. Molecular Pharmacology 72, 103, doi:10.1124/mol.107.035477 (2007).

21 Blechman, J. & Levkowitz, G. Alternative Splicing of the Pituitary Adenylate Cyclase-Activating Polypeptide Receptor PAC1: Mechanisms of Fine Tuning of Brain Activity. Front Endocrinol (Lausanne) 4, 55, doi:10.3389/fendo.2013.00055 (2013).

22 Fradinger, E. A., Tello, J. A., Rivier, J. E. & Sherwood, N. M. Characterization of four receptor cDNAs: PAC1, VPAC1, a novel PAC1 and a partial GHRH in zebrafish. Mol Cell Endocrinol 231, 49–63, doi:10.1016/j.mce.2004.12.002 (2005).

23 Lutz, E. M. et al. Characterization of novel splice variants of the PAC1 receptor in human neuroblastoma cells: Consequences for signaling by VIP and PACAP. Molecular and Cellular Neuroscience 31, 193–209, doi:https://doi.org/10.1016/j.mcn.2005.09.008 (2006).

24 Zhou, C. J. et al. Cellular distribution of the splice variants of the receptor for pituitary adenylate cyclase-activating polypeptide (PAC1-R) in the rat brain by in situ RT-PCR. Molecular Brain Research 75, 150–158, doi:https://doi.org/10.1016/S0169-328X(99)00300-9 (2000).

25 Spongier, D. et al. Differential signal transduction by five splice variants of the PACAP receptor. Nature 365, 170–175, doi:10.1038/365170a0 (1993).

26 Biran, J., Blechman, J., Wircer, E. & Levkowitz, G. in Model animals in neuroendocrinology: From worm to mouse to man (eds Mike Ludwig & Gil Levkowitz) Ch. 5, 101-131 (Wiley-Blackwell, 2018).

27 Steenbergen, P. J., Richardson, M. K. & Champagne, D. L. Patterns of avoidance behaviours in the light/dark preference test in young juvenile zebrafish: A pharmacological study. Behavioural Brain Research 222, 15–25, doi:https://doi.org/10.1016/j.bbr.2011.03.025 (2011).

28 Peng, X. et al. Anxiety-related behavioral responses of pentylenetetrazole-treated zebrafish larvae to light-dark transitions. Pharmacology Biochemistry and Behavior 145, 55–65, doi:10.1016/j.pbb.2016.03.010 (2016).

29 Elbaz, I., Yelin-Bekerman, L., Nicenboim, J., Vatine, G. & Appelbaum, L. Genetic ablation of hypocretin neurons alters behavioral state transitions in zebrafish. The Journal of neuroscience : the official journal of the Society for Neuroscience 32, 12961–12972, doi:10.1523/JNEUROSCI.1284-12.2012 (2012).

30 Maximino, C. et al. Fingerprinting of psychoactive drugs in zebrafish anxiety-like behaviors. PLoS One 9, e103943, doi:10.1371/journal.pone.0103943 (2014).

31 Tran, S. & Gerlai, T. R. The Novel Tank Test: Handling Stress and the Context Specific Psychopharmacology of Anxiety. Current Psychopharmacology 5, 169–179, doi:10.2174/2211556005666160519144414 (2016).

32 Egan, R. J. et al. Understanding behavioral and physiological phenotypes of stress and anxiety in zebrafish. Behav Brain Res 205, 38–44, doi:10.1016/j.bbr.2009.06.022 (2009).

33 Godwin, J., Sawyer, S., Perrin, F., Oxendine, S. E. & Kezios, Z. D. in Zebrafish Protocols for Neurobehavioral Research (eds Allan V. Kalueff & Adam Michael Stewart) 181–189 (Humana Press, 2012).

34 King, S. B., Toufexis, D. J. & Hammack, S. E. Pituitary adenylate cyclase activating polypeptide (PACAP), stress, and sex hormones. Stress 20, 465–475, doi:10.1080/10253890.2017.1336535 (2017).

35 Dias, B. G. & Ressler, K. J. PACAP and the PAC1 Receptor in Post-Traumatic Stress Disorder. Neuropsychopharmacology 38, 245, doi:10.1038/npp.2012.147 (2012).

36 Blechman, J. et al. Specification of hypothalamic neurons by dual regulation of the homeodomain protein Orthopedia. Development 134, 4417–4426, doi:10.1242/dev.011262 (2007).

37 May, V. & Parsons, R. L. G Protein-Coupled Receptor Endosomal Signaling and Regulation of Neuronal Excitability and Stress Responses: Signaling Options and Lessons From the PAC1 Receptor. Journal of Cellular Physiology 232, 698–706, doi:10.1002/jcp.25615 (2017).

38 Spengler, D. et al. Differential signal transduction by five splice variants of the PACAP receptor. Nature 365, 170–175, doi:10.1038/365170a0 (1993).

39 DiCicco-Bloom, E. et al. Autocrine expression and ontogenetic functions of the PACAP ligand/receptor system during sympathetic development. Dev Biol 219, 197–213, doi:10.1006/dbio.2000.9604 (2000).

40 May, V. et al. Pituitary adenylate cyclase-activating polypeptide (PACAP)/PAC1HOP1 receptor activation coordinates multiple neurotrophic signaling pathways: Akt activation through phosphatidylinositol 3-kinase gamma and vesicle endocytosis for neuronal survival. J Biol Chem 285, 9749–9761, doi:10.1074/jbc.M109.043117 (2010).

41 Nicot, A. & DiCicco-Bloom, E. Regulation of neuroblast mitosis is determined by PACAP receptor isoform expression. Proc Natl Acad Sci U S A 98, 4758–4763, doi:10.1073/pnas.071465398 (2001).

42 Ronaldson, E. et al. Specific interaction between the hop1 intracellular loop 3 domain of the human PAC(1) receptor and ARF. Regul Pept 109, 193–198, doi:10.1016/s0167-0115(02)00204-5 (2002).

43 Mustafa, T., Grimaldi, M. & Eiden, L. E. The hop cassette of the PAC1 receptor confers coupling to Ca2+ elevation required for pituitary adenylate cyclase-activating polypeptide-evoked neurosecretion. J Biol Chem 282, 8079–8091, doi:10.1074/jbc.M609638200 (2007).

44 Mustafa, T., Walsh, J., Grimaldi, M. & Eiden, L. E. PAC1hop receptor activation facilitates catecholamine secretion selectively through 2-APB-sensitive Ca(2+) channels in PC12 cells. Cellular signalling 22, 1420–1426, doi:10.1016/j.cellsig.2010.05.005 (2010).

45 Smith, C. B. & Eiden, L. E. Is PACAP the major neurotransmitter for stress transduction at the adrenomedullary synapse? J Mol Neurosci 48, 403–412, doi:10.1007/s12031-012-9749-x (2012).

46 Taupenot, L., Mahata, M., Mahata, S. K. & O’Connor, D. T. Time-dependent effects of the neuropeptide PACAP on catecholamine secretion : stimulation and desensitization. Hypertension 34, 1152–1162, doi:10.1161/01.hyp.34.5.1152 (1999).

47 Donthamsetti, P. et al. Arrestin recruitment to dopamine D2 receptor mediates locomotion but not incentive motivation. Molecular psychiatry, 10.1038/s41380-41018-40212-41384, doi:10.1038/s41380-018-0212-4 (2018).

48 Lesniak, A. et al. Divergent Response to Cannabinoid Receptor Stimulation in High and Low Stress-Induced Analgesia Mouse Lines Is Associated with Differential G-Protein Activation. Neuroscience 404, 246–258, doi:https://doi.org/10.1016/j.neuroscience.2019.02.015 (2019).

49 Usiello, A. et al. Distinct functions of the two isoforms of dopamine D2 receptors. Nature 408, 199–203, doi:10.1038/35041572 (2000).

50 Vagena, E. et al. A high-fat diet promotes depression-like behavior in mice by suppressing hypothalamic PKA signaling. Translational Psychiatry 9, 141, doi:10.1038/s41398-019-0470-1 (2019).

51 Bunea, I. M., Szentágotai-Tătar, A. & Miu, A. C. Early-life adversity and cortisol response to social stress: a meta-analysis. Translational Psychiatry 7, 1274, doi:10.1038/s41398-017-0032-3 (2017).

52 Syed Sheriff, R., Van Hooff, M., Malhi, G., Grace, B. & McFarlane, A. Associations Among Childhood Trauma, Childhood Mental Disorders, and Past-Year Posttraumatic Stress Disorder in Military and Civilian Men. Journal of Traumatic Stress 32, 712–723, doi:10.1002/jts.22450 (2019).

53 Levine, S. Infantile Experience and Resistance to Physiological Stress. Science 126, 405, doi:10.1126/science.126.3270.405 (1957).

54 Lyons, D. M., Parker, K. J. & Schatzberg, A. F. Animal models of early life stress: Implications for understanding resilience. Developmental Psychobiology 52, 402–410, doi:10.1002/dev.20429 (2010).

55 Murthy, S. & Gould, E. Early Life Stress in Rodents: Animal Models of Illness or Resilience? Frontiers in Behavioral Neuroscience 12, doi:10.3389/fnbeh.2018.00157 (2018).

56 Cooper, A. J., Narasimhan, S., Rickels, K. & Lohoff, F. W. Genetic polymorphisms in the PACAP and PAC1 receptor genes and treatment response to venlafaxine XR in generalized anxiety disorder. Psychiatry research 210, 1299–1300, doi:10.1016/j.psychres.2013.07.038 (2013).

57 Han, P. et al. Association of pituitary adenylate cyclase-activating polypeptide with cognitive decline in mild cognitive impairment due to Alzheimer disease. JAMA neurology 72, 333–339, doi:10.1001/jamaneurol.2014.3625 (2015).

58 Van, C. et al. PACAP/PAC1 Regulation of Inflammation via Catecholaminergic Neurons in a Model of Multiple Sclerosis. J Mol Neurosci 68, 439–451, doi:10.1007/s12031-018-1137-8 (2019).

59 King, S. B. et al. The Effects of Prior Stress on Anxiety-Like Responding to Intra-BNST Pituitary Adenylate Cyclase Activating Polypeptide in Male and Female Rats. Neuropsychopharmacology 42, 1679, doi:10.1038/npp.2017.16 (2017).

60 Yan, Y. et al. Pro- and Anti-Mitogenic Actions of Pituitary Adenylate Cyclase-Activating Polypeptide in Developing Cerebral Cortex: Potential Mediation by Developmental Switch of PAC1 Receptor mRNA Isoforms. The Journal of Neuroscience 33, 3865, doi:10.1523/JNEUROSCI.1062-12.2013 (2013).

61 Apostolakis, E. M., Riherd, D. N. & O’Malley, B. W. PAC1 Receptors Mediate Pituitary Adenylate Cyclase-Activating Polypeptide- and Progesterone-Facilitated Receptivity in Female Rats. Molecular Endocrinology 19, 2798–2811, doi:10.1210/me.2004-0387 (2005).

62 Apostolakis, E. M., Lanz, R. & O’Malley, B. W. Pituitary adenylate cyclase-activating peptide: a pivotal modulator of steroid-induced reproductive behavior in female rodents. Mol Endocrinol 18, 173–183, doi:10.1210/me.2002-0386 (2004).

63 Gagnon, J. A. et al. Efficient mutagenesis by Cas9 protein-mediated oligonucleotide insertion and large-scale assessment of single-guide RNAs. PLoS ONE 9, doi:10.1371/journal.pone.0098186 (2014).

64 Blechman, J., Anbalagan, S., Matthews, G. G. & Levkowitz, G. Genome Editing Reveals Idiosyncrasy of CNGA2 Ion Channel-Directed Antibody Immunoreactivity Toward Oxytocin. Frontiers in cell and developmental biology 6, 117, doi:10.3389/fcell.2018.00117 (2018).

65 Montague, T. G., Cruz, J. M., Gagnon, J. A., Church, G. M. & Valen, E. CHOPCHOP: a CRISPR/Cas9 and TALEN web tool for genome editing. Nucleic Acids Research 42, W401–W407, doi:10.1093/nar/gku410 (2014).

66 Borovski, T. et al. Historical and recent reductions in genetic variation of the Sarotherodon galilaeus population in the Sea of Galilee. Conservation Genetics 19, 1323–1333, doi:10.1007/s10592-018-1102-7 (2018).

67 Wircer, E. et al. Homeodomain protein Otp affects developmental neuropeptide switching in oxytocin neurons associated with a long-term effect on social behavior. eLife 6, e22170, doi:10.7554/eLife.22170 (2017).

